# Strength in numbers: collaborative science for new experimental model systems

**DOI:** 10.1101/308304

**Authors:** Ross F. Waller, Phillip A. Cleves, Maria Rubio-Brotons, April Woods, Sara J. Bender, Virginia Edgcomb, Eric R. Gann, Adam C. Jones, Leonid Teytelman, Peter von Dassow, Steven W. Wilhelm, Jackie L. Collier

**Affiliations:** Department of Biochemistry, University of Cambridge, Cambridge, UK; Department of Genetics, Stanford University School of Medicine, Stanford, CA, USA; Institut de Biologia Evolutiva (CSIC-Universitat Pompeu Fabra), Barcelona, Catalonia, Spain; Environmental Biotechnology Lab, Moss Landing Marine Laboratories, CA, USA; Gordon and Betty Moore Foundation, Palo Alto, CA, USA; Department of Geology and Geophysics, Woods Hole Oceanographic Institution, Woods Hole, MA, USA; Department of Microbiology, The University of Tennessee, Knoxville, TN, USA; protocols.io, Berkeley, CA, USA; Facultad de Ciencias Biológicas, Pontificia Universidad Católica de Chile, Santiago, Chile; Instituto Milenio de Oceanografía, Concepción, Chile; School of Marine and Atmospheric Sciences, Stony Brook University, Stony Brook, NY, USA

## Abstract

Our current understanding of biology is heavily based on the contributions from a small number of genetically tractable model organisms. Most eukaryotic phyla lack such experimental models, and this limits our ability to explore the molecular mechanisms that ultimately define their biology, ecology, and diversity. In particular, marine protists suffer from a paucity of model organisms despite playing critical roles in global nutrient cycles, food webs, and climate. To address this deficit, an initiative was launched in 2015 to foster development of ecologically and taxonomically diverse marine protist genetic models. This multifaceted, complex but important challenge required a highly collaborative community-based approach. Herein we describe this approach, the advances achieved, and the lessons learned by participants in this novel community-based model for research.

## I. Introduction and Experimental Model Systems (EMS) Overview

Genetically tractable model organisms have enabled enormous progress in understanding life, but the small number of model organisms available represent a taxonomically restricted view of biological diversity [1,2]. This imbalance means that a relatively small number of organisms have been used to define ‘the canonical’ in eukaryotes. However, eukaryotes are incredibly diverse, with most of this diversity represented by single-celled protists which are typically more phylogenetically divergent from one another than animals are from fungi (Figure 1) [3]. The vast evolutionary history of protists is often reflected in striking differences in gene content, form, behavior, and ecological function [4]. This diversity is central to global biogeochemistry as marine protists drive vital ecological processes, notably marine primary production, carbon capture, and nutrient cycling [5]. In addition, they influence weather cycles, coastal protection by corals, and create population blooms spanning 100s of kilometers with both positive and negative impacts on food production and water quality [6–8]. Thus, understanding the ecological functions of specific marine protists is essential to the understanding of global ecosystems.

**Figure 1.**
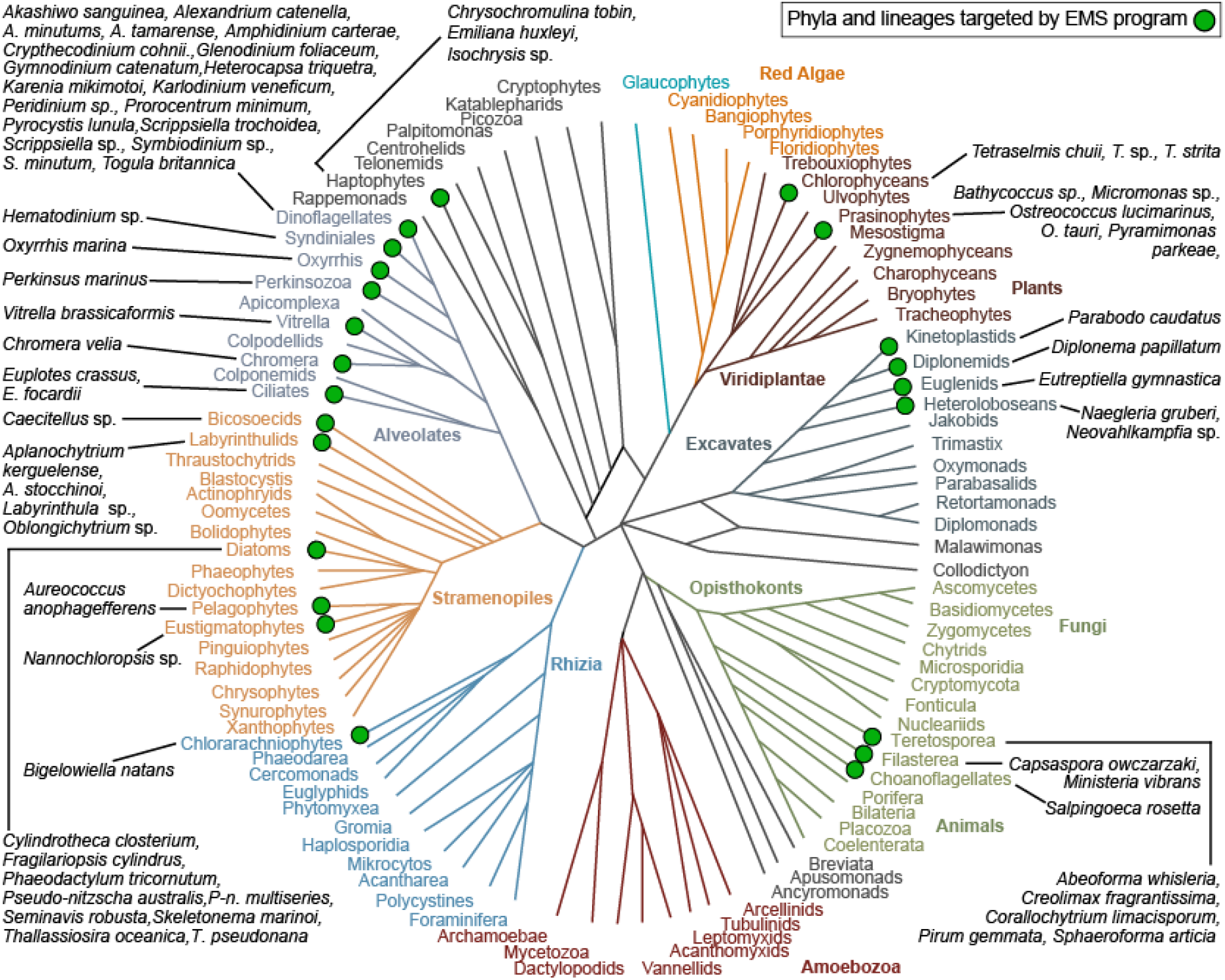
Schematic of relationships of major eukaryotic lineages with taxa the subjects of EMS projects indicated with green dots and listed in black text. Phylogeny modelled on Keeling et al. (2014) [15]

In recent years, genome-wide sequencing technologies have allowed rapid advancement in assessing marine protist genetic diversity1, which has enabled many predictions of cell and organism function based on gene content [9]. These studies have also revealed that often ∼40-60% of the genes in any newly sequenced lineage encode proteins with no known function based on homology with other organisms. For example, about half the genes in a global *Tara* Oceans metatranscriptome assembly have no similarity to known proteins [4]. These unknown genes are likely important in defining distinct ecological contributions of each lineage, however, experiments to determine the functions and importance of such genes are impeded by the lack of genetic methods.

While the need to develop more model marine organisms is clear, many factors create obstacles or disincentives to embracing this challenge. Multiple layers of uncertainty translate to a perceived or actual high risk of failure: Can DNA or RNA be introduced into the relevant cell compartment, and will it be maintained? Will genes be expressed, will their products be detectable, and can selection of transformants be achieved? Will the organism tolerate any or all of these treatments? Furthermore, funding, publishing, and career advancement structures often do not encourage risk-taking enterprises. These compounded uncertainties and risks contribute to a reluctance to commit time and resources to novel genetic model development.

To address these obstacles, in 2015 the Gordon and Betty Moore Foundation launched an $8.1M portfolio of one- and two-year grants as part of its Marine Microbiology Initiative. This new Experimental Model Systems (EMS) program was designed to foster the expansion of genetically tractable models within marine protists, and to promote the power of genetic approaches for studying marine microbial processes amongst the scientists and students working in these fields. Based on early success of the program, many projects were continued through an additional $4.5M of support in 2017.

Beginning with an open call for proposals, the Moore Foundation selected 34 project teams to develop new genetic tools in marine protists. Teams were encouraged to be interdisciplinary—often featuring complementary experts with experience in well-established model systems, such as yeast, as well as marine protists—providing a mechanism to bring diverse skills and innovative approaches to the projects. Nine of the EMS projects focused on protists for which basic genetic techniques were already available (primarily diatoms) enabling the development of more advanced forward and reverse genetics methods. The remaining 25 projects focused on a diverse array of protists for which reliable transformation had not yet been achieved. This project portfolio had the explicit goal of enabling the scientists working on less-explored organisms to learn directly from the successes made with the more advanced model protists. A key EMS strategy was to provide strong support for community building and dialogue between groups to encourage productive interactions and sharing. Over the course of the two years, this EMS effort developed a novel model for research that has enabled significant progress in genetic tool development in marine protists. This model is broadly applicable to other groups of scientists and/or funding organizations wanting to embark on equivalent high-risk complex projects, such as the US National Science Foundation’s <u>E</u>nabling <u>D</u>iscovery through <u>GE</u>nomic Tools (EDGE) program.

## II. Organisms and Approaches

Target organisms were selected to represent the broad ecological and phylogenetic diversity of marine protists. Most of the major eukaryotic super-groups were represented: opisthokonts, excavates, viridiplantae, alveolates, stramenopiles, haptophytes, and rhizarians (Figure 1). Nearly half of the targeted organisms were photoautotrophs, ∼40% were osmoheterotrophs or phagotrophs, and the remainder were mixotrophs (both phototrophic and heterotrophic) or parasitic. Most of the targeted organisms were from temperate ocean habitats, although three polar organisms were included. Half of the targeted organisms were planktonic including species capable of forming harmful algal blooms (mainly dinoflagellate and diatom), and approximately 20% of the targeted organisms were symbionts, including parasites, of marine plants or animals. Although most protists studied were available in existing laboratory cultures, a small number of projects also attempted to directly transform natural protist communities within environmental samples. The broad assortment of growth requirements, cell surface and wall structures, genome organizations, and gene regulation mechanisms of the target protists posed a multifaceted challenge to the community engaged in developing these new genetic model organisms. However, this diversity also provided an opportunity to investigate the mechanistic basis of a wide variety of biological processes in the ocean.

After the first two years of the EMS program, EMS researchers were surveyed to capture the breadth of taxa used and transformation techniques trialed, while also measuring successes achieved and persistent challenges. At least seven different methods of DNA or RNA introduction were attempted (Table 1), with electroporation being the method both most tested and successful. The strongest collective effort was applied to diatoms (Bacillariophyceae) and ‘core’ dinoflagellates (Dinophyceae) owing to their abundance, diversity, and ecological importance [10]. The transgenes used were typically for expression of antibiotic resistance proteins for selection, or fluorescent reporter proteins for visual identification of successful transformants (Table 1). In most cases, these transgenes were flanked by native promoter and terminator elements to increase the likelihood of expression. Other strategies used ‘universal’ promoters (e.g., 35S and CMV) when targeting multiple diverse taxa simultaneously, including two efforts to bulk transform natural plankton communities. Of the 15 major taxonomic classes represented in the survey results, stable transformation was achieved in six and at least transient expression of the transgene was achieved in an additional six (Table 1, Figure 2). Survey respondents rated how much effort they had put into each transformation method and taxon as: high, medium, or low. The amount of effort generally positively correlated with at least some success of transgene expression and/or maintenance, providing evidence for the value of long-term support for such ventures. The taxonomic group most recalcitrant to transformation despite effort was the dinoflagellates, a group whose distinctive nuclear biology is also least well understood [11–13].

**Table 1:**
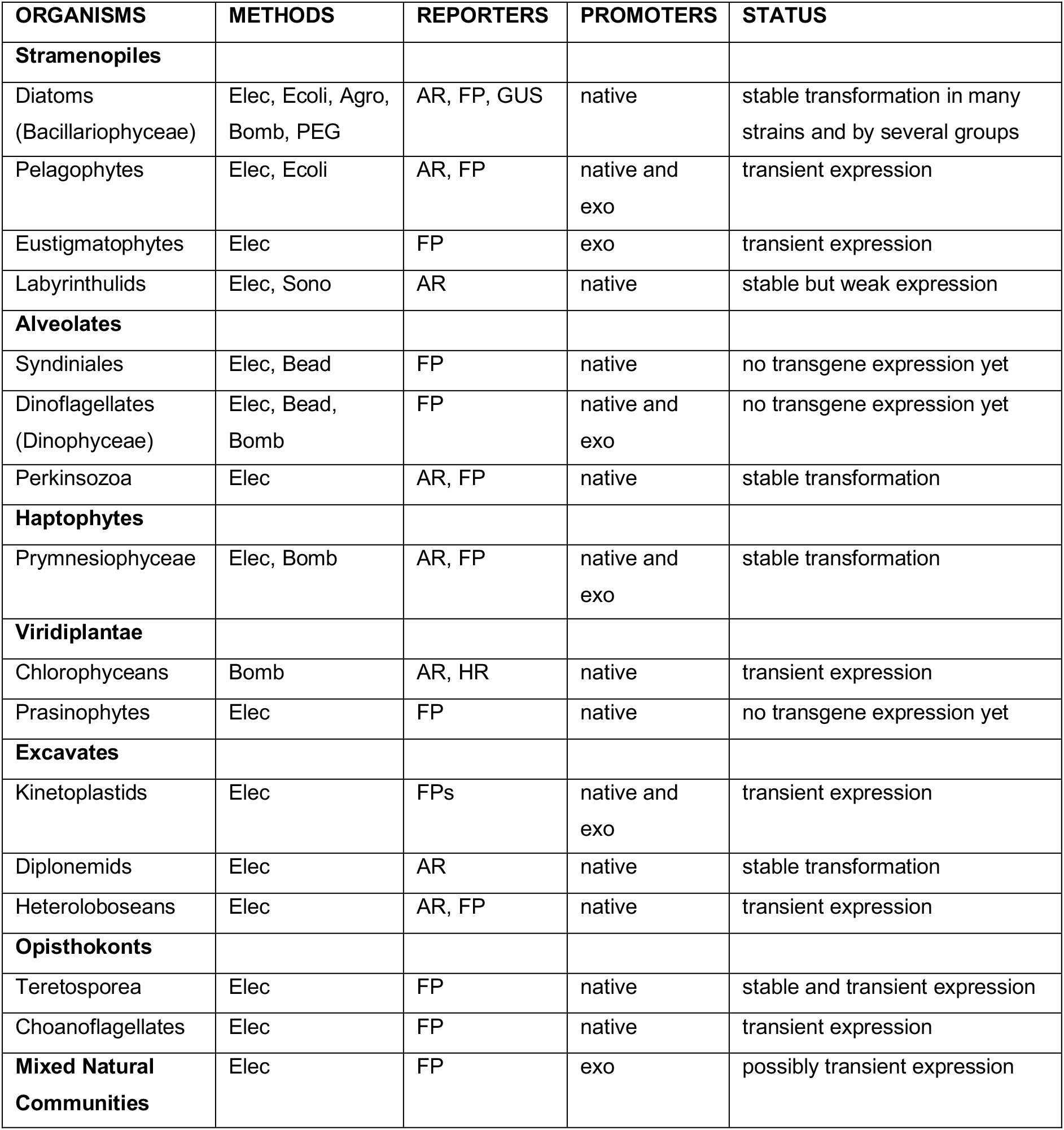
EMS investigators’ survey summary. Methods of transfection: Elec; electroporation: Ecoli; conjugation with *E. coli*: Agro; conjugation with *Agrobacterium*: Bomb; particle bombardment: PEG; Polyethylene glycol: Sono; sonoporation: Beads; glass beads. Reporters: AR; antibiotic resistance: HR; herbicide resistance: FP; fluorescent proteins: GUS; beta-glucuronidase. Promoters: transgenes driven by native or exogenous; exo promoters.

**Figure 2.**
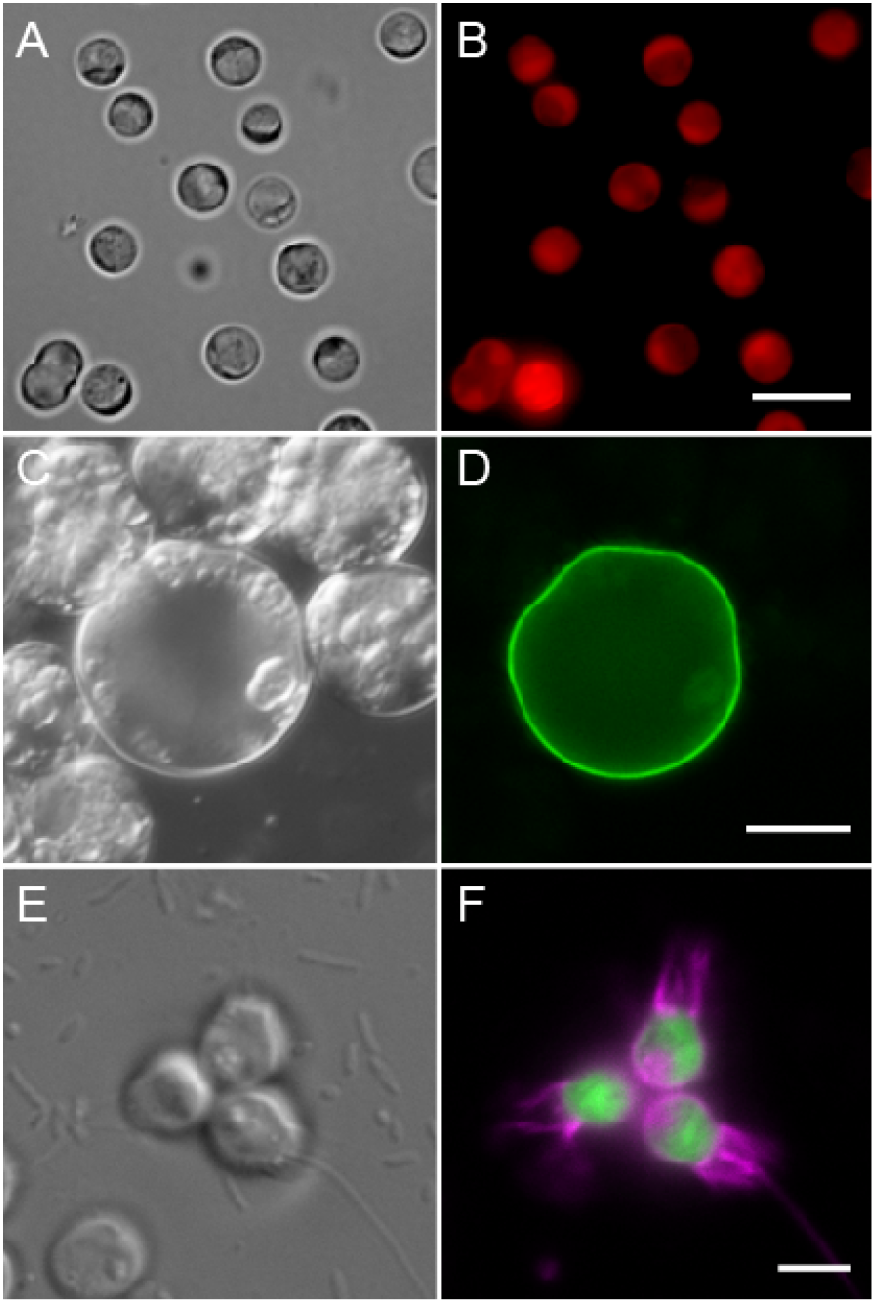
Examples of EMS transformed protists. A,B) *Corallochytrium limacisporum* stably expressing the mCherry fluorescent protein (red) fused to a puromycin resistance protein driven by an endogenous actin promoter (M. Rubio-Brotons, UPF-CSIC, Barcelona, Spain). C,D) *Perkinsus olseni* (marine bivalve parasite) expressing a GFP (green) fusion with an exported cell wall protein (R. Waller, University of Cambridge, UK). E,F) The choanoflagellate *Salpingoeca rosetta* transformed with a plasmid expressing fluorescent proteins that illuminate the cell body (green) and the plasma membrane (magenta). (D. Booth, U. California Berkeley, USA). Scale bar = 5 μm, 20 μm and 5 μm for B, D, F respectively.

## III. Building Community

A key component of the EMS strategy was to support exchange and discussion of methods details among EMS researchers. To achieve this goal, Moore Foundation staff and research teams engaged in multiple approaches to foster communication and collaboration. The intent was to create an environment open to sharing and with a low barrier of entry, to disseminate both positive as well as negative results rather than only the positive outcomes typically released via peer-reviewed publication, and to encourage rapid sharing in weeks or months rather than years. Early in the program cycle a ‘kick-off’ meeting was held in Heidelberg, Germany, including approximately 50 of the scientists involved and across all career stages. This meeting focused on building connections and sharing ideas and strategies. Several further Foundation-sponsored EMS-themed conference events were held throughout the grant period to encourage further interaction both between teams and beyond. In parallel with these in-person meetings, quarterly ‘Virtual Convening’ webinars provided a forum for all community members to share the progress and challenges of their projects. These events featured two or three speakers working on different taxa and ample time for questions and feedback from the community, with 25-30 participants per convening. The virtual gatherings brought together researchers in a ‘lab meeting’ atmosphere without requiring additional travel and resources.

To further accelerate the pace of idea exchange among researchers, the EMS program supported the web-based open-access platform *protocols.io* [14] to establish a dedicated user group, ‘Protist Research to Optimize Tools in Genetics’ (PROT-G). This online group facilitated protocol sharing and discussions among the grantees and any other researchers interested in protist genetics. *Protocols.io* staff enlisted graduate student and postdoctoral site users to act as ambassadors to promote PROT-G at their institutions and at conferences. As of April 2018, PROT-G contained 139 public protocols, 152 members, and 44 ongoing discussions that map globally between EMS research teams (Figure 3). Survey data from the EMS community also reported that most respondents (∼80%) had engaged in and benefited from interactions between teams that the virtual meetings and PROT-G network had allowed, including the sharing of ‘negative’ results (Figure 3). This community-oriented model of science operating independently from the traditional publication pathway has very clear benefits for tackling challenging, multifaceted research programs.

**Figure 3.**
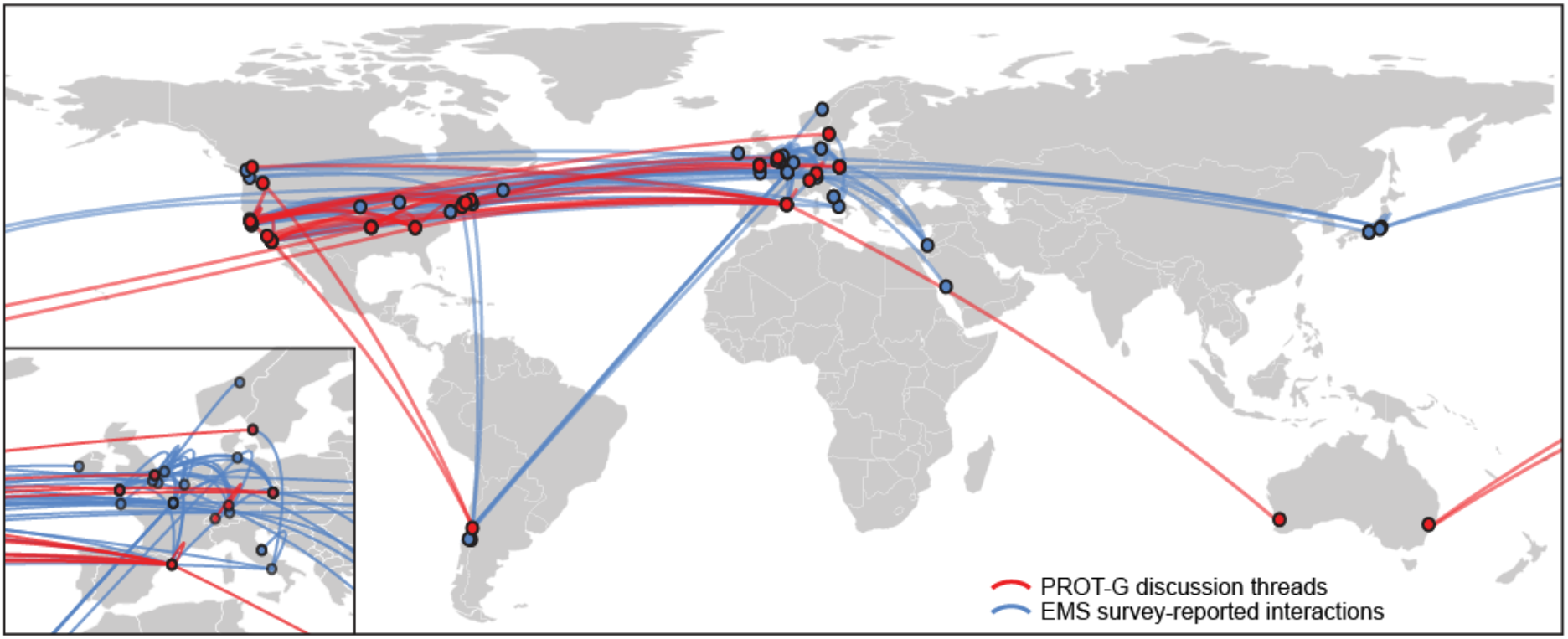
Global EMS network map of protocols.io PROT-G discussion threads (red) and further direct discussions and interactions (blue) between program teams reported in the EMS survey.

## IV. Lessons Learned, Challenges, and the Future

The prospect of failure often deters scientists from embarking on high-risk projects. ‘Negative’ results are viewed as difficult to publish, and are frequently buried in laboratory notebooks with, at best, some word-of-mouth communication between colleagues. The value of sharing these results, and making such information readily accessible, was brought into clear focus through the frequent meetings and discussions within the EMS community. Our concerted effort to genetically transform new and often recalcitrant taxa provided a platform for discussion and comparison of approaches taken enabling a fuller assessment of different methods, progress, and outstanding challenges. *Protocols.io* provided a platform for this sharing of methods, irrespective of their level of progress towards success, and these published methods now provide an inventory of details of approaches taken, effort made, outcomes as they stand, and contact details of the researchers involved. This evolving resource will continue to inform and support researchers engaging in the EMS challenge.

The program recognized from the outset that this large, multivariate challenge required broad and cooperative contributions from many research teams representing different expertise and experience. A community was built that nucleated around a set of common goals, minimized effort duplication and leveraged the research strengths of each team. For many program members this community-based effort was a unique research experience, and a welcome alternative to individual, competitive-style research. An initial challenge was to overcome the tendency of scientific teams to keep research results private until publication. This challenge was met by the early use of the Virtual Convening events to share initial strategies, data of transformation attempts and progress updates, making the benefits of this exchange quickly apparent. A sense of community participation and support was also fostered by researchers publishing and discussing experimental method details in the protocols.io domain before release into the peer-reviewed public space. These strategies for community formation and participation are likely relevant to many other multi-group research programs.

From a funding perspective, the EMS strategy was to spread resources among a relatively large number of teams working on different organisms and approaches. One year of funding was provided to a wide collection of teams to ‘test the water’ for early signs of organism genetic tractability. Funding was then provided, for up to two additional years, where early progress was demonstrated. This element of the program aimed to maximize the likelihood for positive outcomes and to lower risk for overall progress where success was otherwise difficult to predict among such diverse biological systems. While this provided welcome support for high-risk endeavors, it remains difficult to assess if the persistent challenges in some taxa would be overcome with more time and greater effort.

The EMS program has ignited new drive and progress to overcome what many in the fields of marine protistology, ecology, and oceanography have recognized as a significant obstacle to understanding these complex and important biological systems. But how do researchers maintain this momentum beyond the duration of the EMS program? For the organisms still recalcitrant to transformation, how can scientists move forward avoiding the notion that this cause would not benefit from additional support? These are some of the challenges that the EMS community now must grapple with and embrace. For many involved, however, this program has also been a positive new experience in community-driven research. It has provided a model for tracking, quantifying and sharing progress across a wide network much more rapidly and completely than the traditional publishing paradigm. Furthermore, there has been real collective success in technology development and diffusion (including 17 publications citing EMS funding as of April 2018), which has substantially lowered the barriers to testing new organisms as potential future models. As individual teams’ research programs now move forward we will see the rewards of hypothesis-driven experiments that these new tools allow, and advances in our understanding of ocean organisms and ecosystems. Applying this momentum to surmounting other large, complex obstacles will require continued recognition and promotion of the value of ongoing support for technology development programs such as EMS both to federal funding agencies and policy makers, as well as to charitable trusts and foundations.

## Acknowledgments

We thank the EMS Advisory Committee: Bonnie Bassler (Princeton), Vicki Chandler (Minerva Schools), and John Pringle (Stanford), and the many researchers participating in the EMS effort. We also thank Jose-Antonio Fernandez Robledo for discussions on the manuscript. The work described here and the growing connections and collaborations among the authors were supported by the Gordon and Betty Moore Foundation’s Marine Microbiology Initiative.

